# Brain age predicted using graph convolutional neural network explains developmental trajectory in preterm neonates

**DOI:** 10.1101/2021.05.15.444320

**Authors:** Mengting Liu, Sharon Kim, Ben Duffy, Shiyu Yuan, James H. Cole, Arthur W. Toga, Neda Jahanshad, Anthony James Barkovich, Duan Xu, Hosung Kim

**Affiliations:** Department of Neurology, USC Stevens Neuroimaging and Informatics Institute, Keck School of Medicine, University of Southern California, Los Angeles, CA; Centre for Medical Image Computing, Department of Computer Science, University College London, London, UK; Department of Radiology & Biomedical Imaging, University of California, San Francisco, San Francisco, CA

**Keywords:** brain age prediction, Graph convolutional network, neonates, relative brain age, structural equation modeling

## Abstract

Dramatic alterations in brain morphology, such as cortical thickness and sulcal folding, occur during the 3rd trimester of gestation which overlaps with the period of premature births. Here, we investigated the ability of the graph convolutional network (GCN) to predict brain age for preterm neonates by accounting for morphometrics measured on the cortical surface and the surface mesh topology as a sparse graph. Our findings demonstrate that GCN-based age prediction of preterm neonates (n=170; mean absolute error [MAE]: 1.06 weeks) outperformed conventional machine learning algorithms and deep learning methods that did not use topological information. We further evaluated how predicted brain age (PBA) emerges as a biologically meaningful index that characterizes the current status of brain development at the time of imaging. We hypothesized that the relative brain age (RBA; PBA minus chronological age) at scan reflects a combination of perinatal clinical factors, including preterm birth, birthweight, perinatal brain injuries, exposure to postnatal steroids, etc. We also hypothesized that RBA of neonatal scans may be associated with brain functional development in the future. To validate these hypotheses, we used general linear models. Furthermore, we established structural equation models (SEM) to determine the structural relationship between preterm birth (as a latent variable of birthweight and birth age), perinatal injuries (as a latent variable of three leading brain injuries), postnatal factors (as a latent variable of six clinical conditions), RBA at scan, and neurodevelopmental scores at 30 months. Our results suggest that low birthweight, chronic lung disease, and exposure to postnatal steroids impair cortical growth, as low RBA was significantly associated with these risks. Furthermore, RBA was associated with cognitive and language scores at 30 months. SEM analysis indicated that RBA mediated the influences of preterm birth and postnatal clinical factors, but not perinatal brain injuries, toward brain functional development at 30 months. The left middle cingulate cortex showed the most accurate prediction of brain age (MAE: 1.19 weeks), followed by left posterior and right middle cingulate cortices (1.21 weeks). These cingulate regions presented faster growth than others. RBAs of several frontal cortices significantly correlated with cognitive abilities at 30 months of age (n=50). Whereas, RBA of left Broca’s area, which is important for language production and comprehension, was associated with language functional scores. Overall, our results demonstrate the potential of the GCN in both predicting brain age and localizing regional growth that relates to postnatal factors and future neurodevelopmental outcome.

## Introduction

Predicting brain age using neuroimaging data and machine learning approaches has emerged as a biologically meaningful index that assesses the status of brain development or aging at the time of imaging (Jónsson *et al.*, 2019). Particularly, in early development, estimating the age of neonatal brains using the predicted brain age (PBA) measurement may be clinically useful in evaluating neurodevelopment and possible impaired brain growth led by perinatal factors. Indeed, a growing body of evidence has demonstrated that brain structural and functional maturation is impaired in preterm infants, including decreased cerebral volume (Ajayi-Obe *et al.*, 2000; Kapellou *et al.*, 2006), altered cortical surface area and microstructural organization (Ball *et al.*, 2013) as well as aberrant functional and structural connectivity (Pandit *et al.*, 2014; Smyser *et al.*, 2016). Moreover, various perinatal factors, such as birthweight, brain injuries, gestational age, and chronic lung disease of prematurity (CLD), also termed bronchopulmonary dysplasia, have been confirmed to alter brain development and outcomes of preterm neonates (Thompson *et al.*, 2007). Thus, it remains clinically imperative to identify robust metrics that assess how various perinatal factors affect neurodevelopment and outcome. PBA, with its increasing recognition in biological relevance, may provide such clinical utility in assessing preterm brain development throughout the third trimester of gestation.

To study brain aging using PBA, studies to date particularly focused on using machine learning and deep learning (DL) methods (LeCun *et al.*, 2015; Peng *et al.*, 2019) that learn important features without priori information or hypotheses. For instance, in aging brains, the PBA has been investigated using standard deep learning approaches (Ning *et al.*, 2020) where an image volume is inputted into a convolutional neural network (CNN) and a number representing the whole brain age is outputted (Huang *et al.*, 2017). Notably, Cole et al. (Cole *et al.*, 2017) implemented a deep learning model trained by T1-weighted MRIs to predict brain age and achieved promising results in PBA’s association with biological variables of aging. Similar approaches have been executed in neonatal data, but the PBA models have been applied only to connectivity data (Kawahara *et al.*, 2017), structural connectivity data (Brown *et al.*, 2017), and myelin-based brain features (Ouyang *et al.*, 2019), and showed lack of sensitivity in predicting neurodevelopmental outcome (He *et al.*, 2018).

The analysis of specific morphological features may enhance PBA accuracy, given that the third trimester involves highly coordinated morphological alterations (Kim *et al.*, 2020). During the perinatal and the postnatal period, brain morphology undergoes a doubling of whole brain volume (Sherer *et al.*, 2007; Salmaso *et al.*, 2014), a fourfold increase of cortical gray matter volume. By the first postnatal year, cerebral cortical surface area expands approximately 80% and cortical thickness increases about 42% (Li *et al.*, 2015a). Thus, unlike prior approaches relying on deep learning models to output potentially useful features from brain MR images, we specifically focused on highly age-related features within the third trimester, such as increases in cortical folding or volume, to enhance neonatal brain age prediction. However, these morphological properties, including sulcal depth, cortical thickness, and cortical gray matter/superficial white matter intensity ratios (Lewis *et al.*, 2018), are conventionally difficult to characterize through the direct analysis of the voxel-wise MR images, since these features are usually extracted onto a form of manifolds comprising cortical surfaces. Thus, obtaining cortical surfaces by tessellating the brain surface into triangulated brain meshes of vertices (Fischl, 2012) is a prerequisite.

Additionally, a large number of edges linking between the neighboring vertices (i.e., points on the cortical surface) involve important topological information, representing the location and adjacency among neighboring vertices. In this context, conventional CNN methods may not be suited for the analysis of cortical morphological features, and traditional machine learning models applied to cortical features have not taken into account the topological relationship of vertices. Thus, we investigated the ability of a graph convolutional network (GCN) (Defferrard *et al.*, 2016) to predict preterm neonatal brain age when accounting for the mesh topology as a sparse graph using surface-wise cortical morphometrics, i.e., cortical thickness, sulcal depth and gray/white intensity ratio.

PBA presents values older or younger than the true chronological age. While a positive value in relative brain age (RBA; PBA minus chronological age) is indicative of compromised health status in adults (Franke and Gaser, 2019), where RBA has been reported to be positively correlated with neurological diseases and impaired functional outcomes in adult brain studies, a negative RBA in the neonatal cohort may indicate impaired or delayed neurodevelopment. We hypothesize that the RBA at neonatal scan reflects the collective effects of pre-scan (pre- and postnatal) clinical factors, including the severity of preterm birth, perinatal brain injuries, postnatal treatments, neonatal infections, and postnatal cardiorespiratory complications. We furthermore hypothesized that an RBA could be a predictor of neurodevelopmental outcome in preterm survivors. Taking these hypotheses into account, we explored the accuracy and clinical utility of PBA using a graph-based CNN (GCN) model that enables the convolutional filtering of input features through surface topology in the context of spectral graph theory (Shuman *et al.*, 2013). Cortical thickness, sulcal depth, and GM/WM intensity ratio maps were extracted from the cortical mesh and inputted to the GCN. To evaluate the GCN-based PBA, we assessed: 1) whether the GCN approach outperforms other machine learning algorithms that do not consider surface topology in learning vertex-wise features and predicting brain ages; 2) whether the PBA using GCN reflects the influence of pre-scan clinical factors on neurodevelopment; 3) whether RBA of neonatal MRI is a sensitive predictor of neurodevelopmental outcome in 30 months for preterm survivors; 4) whether RBA at scan mediates the relationship between preterm birth-related clinical factors and neurodevelopmental behavioral performance at 30 months; and 5) whether the GCN can be expanded to predict regional brain ages (i.e., identifying which regions specifically are affected by injuries; determining whether certain regions mature at different rates relative to other regions).

## Materials and Methods

### Subjects

Our dataset comprised of 129 preterm neonates (mean postmenstrual age at birth [PMA] = 28.2±1.9 weeks; range 24 –33 weeks) admitted to UCSF Benioff Children’s Hospital San Francisco between June 2008 and May 2017 (Table 1). Most subjects were scanned twice, but some scans were excluded due to a large amount of motion artifact, providing a total of 170 MRI scans (PMA at 1^st^ scan: 31.4±1.9 weeks; range 26.7– 35.7 weeks; 2^nd^ scan: 36.0±1.9 weeks; range 32.1–43.4 weeks). Parental consent was obtained for all cases following a protocol approved by the Institutional Committee on Human Research.

**Table 1.**
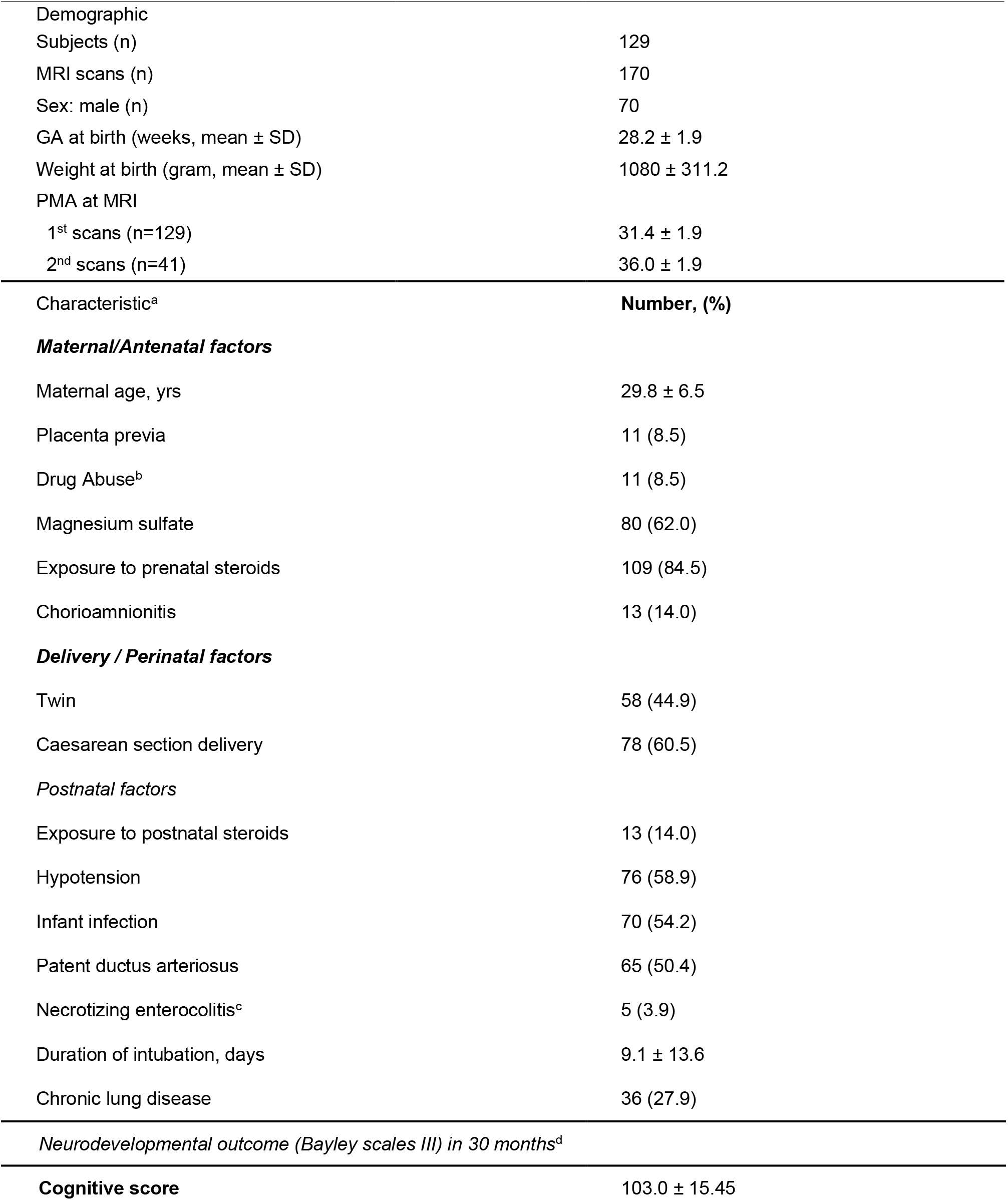
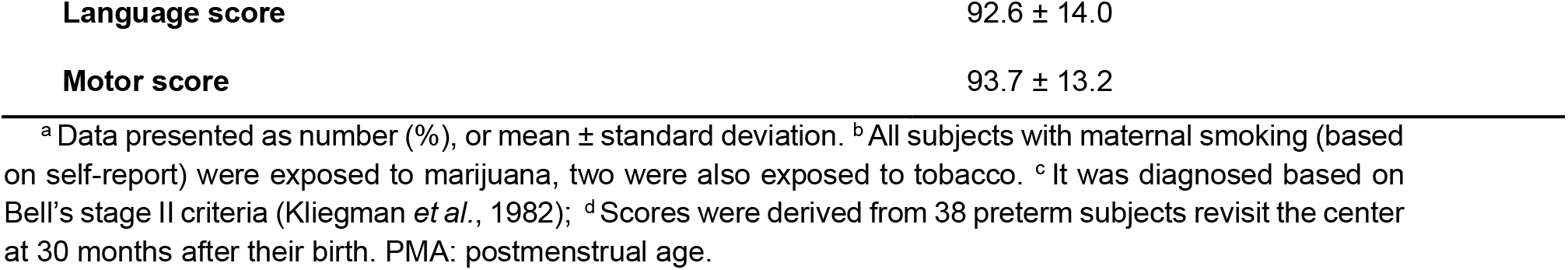
Demographic and clinical characteristics for preterm neonates

### MRI acquisition

Newborns enrolled between 2008 and 2011 (n = 56) were scanned on a 1.5-Tesla General Electric Signa HDxt system using a specialized high-sensitivity neonatal head coil built within a custom-built MRI-compatible incubator. T1-weighted images were acquired using sagittal 3-dimensional inversion recovery spoiled gradient echo (3D SPGR) (TR = 35; TE = 6; FOV = 256×192 mm^2^; number of excitations [NEX] = 1; and FA = 35°), yielding images with 1×1×1 mm^3^ resolution. Newborns enrolled between 2011 and 2017 (n = 73) were scanned on a 3-Tesla General Electric Discovery MR750 system. T1-weighted images were acquired using sagittal 3D IR-SPGR (inversion time of 450 ms; FOV = 180×180 mm^2^; NEX = 1; FA = 15°), yielding images with 0.7×0.7×1 mm^3^ resolution.

### Clinical factors

Neonatal demographic and clinical variables were collected by a trained clinical research nurse (Table 1). Newborns with culture-positive sepsis, clinical signs of sepsis with negative blood culture, or meningitis were classified as having infection. Newborns with clinical signs of patent ductus arteriosus (PDA; prolonged systolic murmur, bounding pulses, and hyperdynamic precordium), and evidence of left-to-right flow through the PDA on echocardiogram were classified as having a PDA. Necrotizing enterocolitis (NEC) was diagnosed according to Bell stage II criteria or higher.

For diagnostic of neonatal brain injuries, a pediatric neuroradiologist (A.J.B.) blinded to patient history reviewed patient MRI scans, including 3-D T1 and axial T2- weighted sequences, as well as SWI when available. Presence and severity of three leading drivers of neurodevelopmental deficits, i.e. intraventricular hemorrhage (IVH), ventriculomegaly (VM), and periventricular leukomalacia (PVL) or white matter injury, were visually scored. The severity scores were generated for IVH using the scoring system of Papile (0: absent; 1: germinal matrix hemorrhage; 2: IVH; 3: IVH with hydrocephalus; 4: parenchymal hemorrhage, usually periventricular hemorrhagic infarction) (Papile *et al.*, 1978) and WMI (0: absent; 1: 2 mm; 3: >5% hemisphere) using established criteria (Papile *et al.*, 1978; Miller *et al.*, 2003). Subsequently, IVH scores were binarized with “mild” representing grades 1– 2, and “severe” representing grades 3–4; WMI and VM were categorized as “mild” for grade 1 and “severe” for grades 2– 3. For subjects with multiple MR examinations, the highest (most severe) score in each category was used for analysis. In the current study, we merged infants with mild injuries and those with no injury into one none-mild injury group since the two groups exhibited no significant differences in the following analyses.

### Neurodevelopmental assessment

All the infants in our analysis were referred to the UCSF Intensive Care Nursery Follow-Up Program upon discharge for routine neurodevelopmental follow-up. Neurodevelopment was assessed using the Bayley-III, which was performed by unblinded clinicians at 30 months’ corrected age. We assessed cognitive, verbal/language and neuromotor performance. Follow-up was available in 38 of the 129 infants who survived to hospital discharge.

### Image processing and cortical surface extraction

The cortical surfaces were constructed using the NEOCIVET pipeline (Kim *et al.*, 2016; Liu *et al.*, 2019; Liu *et al.*, 2021). The pipeline began with general MR image pre-processing, including denoising and intensity nonuniformity correction. Then, the brain is extracted using a patch-based brain extraction algorithm (BEaST) (Eskildsen *et al.*, 2012) and registered to the MNI-NIH neonatal brain template (http://www.bic.mni.mcgill.ca/ServicesAtlases/NIHPD-obj2). Different types of brain tissue (GM, WM, and CSF) were thereafter segmented by a label fusion based on a joint probability between selected templates (Wang *et al.*, 2013). Next, the corpus callosum was segmented on the midline-plane and used to divide the WM into hemispheres. A marching-cube based framework was adopted to generate a triangulated mesh WM surface attached to the boundary between the GM and WM. After resampling to a fixed number of 81,920 surface meshes using the icosahedron spherical fitting, this surface was further fitted to the sharp edge of the GM-WM interface based on the image intensity gradient information. This allowed for the deformation while preserving the spherical topology of the cortical mantle. A CSF skeleton was then generated from the union of WM and CSFs. Pial surface was constructed by expanding the WM surface towards the skeleton as an intermediate pial surface. The intermediate pial surface further underwent a fine deformation to identify actual edges of sulcal CSF volumes using an intensity gradient feature model. Finally, the cortical thickness was estimated based on the Euclidean distance between the white matter and pial surface.

The cortical morphology was quantitatively characterized by measuring cortical thickness, sulcal depth, and gray/white intensity ratio (Lewis *et al.*, 2018) on the cortical surface at 81,924 vertices (163,840 polygons). These features were further re-sampled to the surface template using the transformation obtained in the surface registration, to allow for inter-subject comparisons.

### Graph convolutional neural network (GCN) based brain age prediction

The proposed PBA model using GCN is illustrated in Figure 1. GCNs (Defferrard *et al.*, 2016) are designed to exploit the underlying graph structure of the data. To this end, GCNs consider spectral convolutions on graphs defined as the multiplication of a signal with a filter in the Fourier domain (Shuman *et al.*, 2013). The signal *h* on the graph nodes is filtered by *g* as:

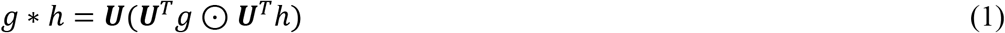

**Figure 1.**
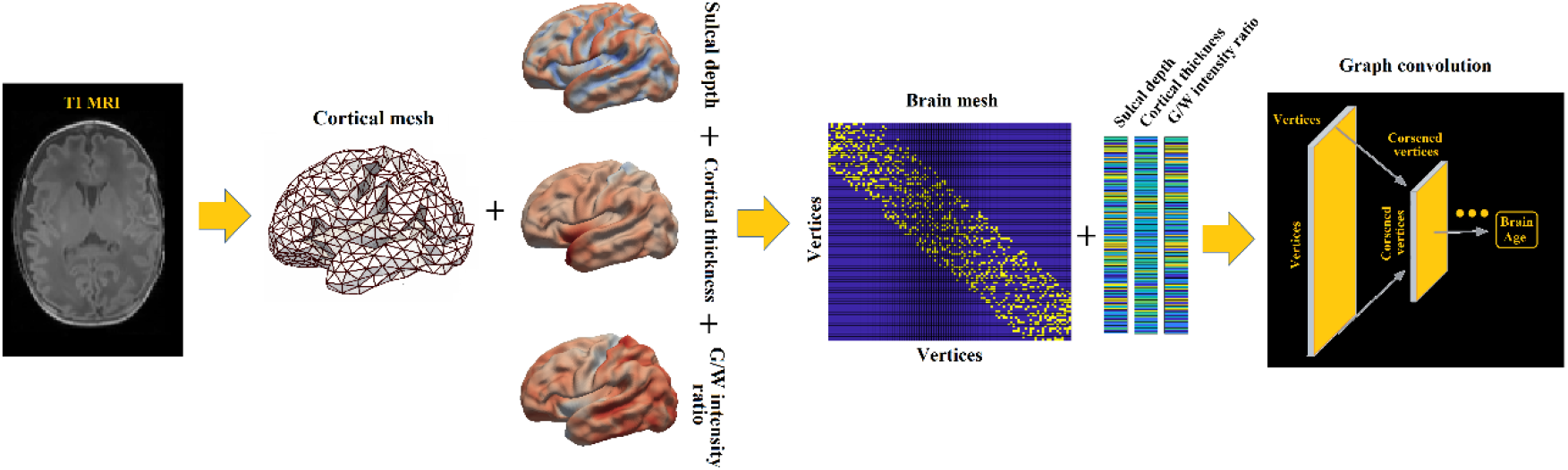
The proposed graph-based convolutional network for brain age prediction.

where U is the Fourier basis of the graph Laplacian L, given by the eigendecomposition of L i.e., **L** = ***U*Λ*U^T^*. Λ** is the ordered real nonnegative eigenvalue values vector of graph Fourier transform, * is the convolution operator and ⊙ denotes element-wise multiplication. The graph Laplacian L is defined as L: = D – W where the degree matrix D is a diagonal matrix whose *i*th diagonal element *d_i_* is equal to the sum of the weights of all the edges connected to vertex i as *d_ii_* = Σ*_j_ W_ij_*. W is a binarized adjacency matrix encoding the connection between vertices, where 1 represents a connection between two vertices in brain mesh and 0 otherwise. After normalization, the graph Laplacian is defined as **L** = *I_n_* **− D**^−1/2^**WD**^−1/2^ where *I_n_* is the identity matrix.

Vertices on graph are re-arranged such that a graph pooling operation becomes as efficient as 1D pooling. Fake nodes, or disconnected nodes, are added to construct a balanced binary tree from the coarsest to finest level to make the pooling operation very efficient without losing information.

We down-sampled 81,924 vertices on cortical surfaces using the icosahedron downsampling to investigate the accuracy of the GCN-based brain age prediction while saving computational time in the training of GCN. We thus feed each of the brain meshes that were down-sampled with 324, 1,284, 5k and 20k vertices to the GCN model respectively. The number of 1,284 was chosen to use in the following analysis by a compromise of the computational accuracy and computational time (Figure S1).

In the current study, the input graphs combined cortical thickness, sulcal depth, and GM/WM intensity ratio mapped on the surface meshes with an adjacency matrix representing the mesh topology. The feature maps were inputted as vector valued signals on the graph nodes and the sparse binary adjacency matrix by which the connections between each vertex and its neighbor vertices are defined. Mean squared error (MSE) was used as the loss function with an Adam optimizer, the empirically determined set of parameters with a learning rate of 10^−6^, an L2 regularization parameter of 10^−8^, and a batch size of 2 were applied.

### Relative brain age

After calculating predicted brain age for each subject, we further calculated a metric that reflected a subject’s relative brain health status, called relative brain age (RBA). RBA was initially measured by subtracting true brain age from predicted brain age (Cole *et al.*, 2017). Due to regression dilution (Ning *et al.*, 2020), however, it is also possible that regression models bias the predicted brain age toward the mean, underestimating the age of older infants and overestimating the age of younger infants (Brown *et al.*, 2017). When deriving the RBA, this bias thus needs to be corrected using a strategy that was introduced in (Smith *et al.*, 2019; Ning *et al.*, 2020). We defined the new RBA as the difference between the individual RBA and the expected RBA (measurement fitted over the entire sample set by the regression model and leave-one-out cross-validation). The RBA was corrected in this way such that the RBAs of the whole dataset analyzed became unbiased across all age ranges (Figure S2).

To validate our hypothesis that perinatal clinical factors negatively affect brain growth (lower RBA), each variable was dichotomized using clinically defined categorization or median if arbitrary (Table S1). Necrotizing enterocolitis (NEC) was not included in the analysis due to the very small number of subjects (n=5). We then tested the group difference in RBA for each variable separately in a univariate fashion while correcting for postmenstrual age (PMA) at scan and other clinical factors. To this end, we used a general linear mixed-effect model that addressed changes of within- and between-subject effects, and to remove the effects from clinical covariates other than the main variable. In this analysis, only images with all clinical information available were kept (n = 121).

### Structural Equation Modelling

We built structural equation models (SEM) that impute relationships between latent variables. Based on the hypothesized latent risk variables and timeline in Figure 2, we analyzed multiple relationships/paths between severity of preterm birth (as a latent variable of birthweight and birth age), perinatal injuries (as a latent variable of three main types of brain injuries related to preterm birth), pre-scan post-natal factors (as a latent variable of 6 clinical conditions/procedures), RBA at postnatal scan (as a manifest variable) and neurodevelopmental outcome scores at 30 months (as a latent variable of 3 brain functional scores). Fifty neonatal MR images including baseline and follow-up scans were available from 38 preterm survivors who revisited the center 30 months. This analysis, which was designed to identify the clinical variables and their paths leading to adverse neurodevelopmental outcomes, was conducted on the 50 MRI images.

**Figure 2.**
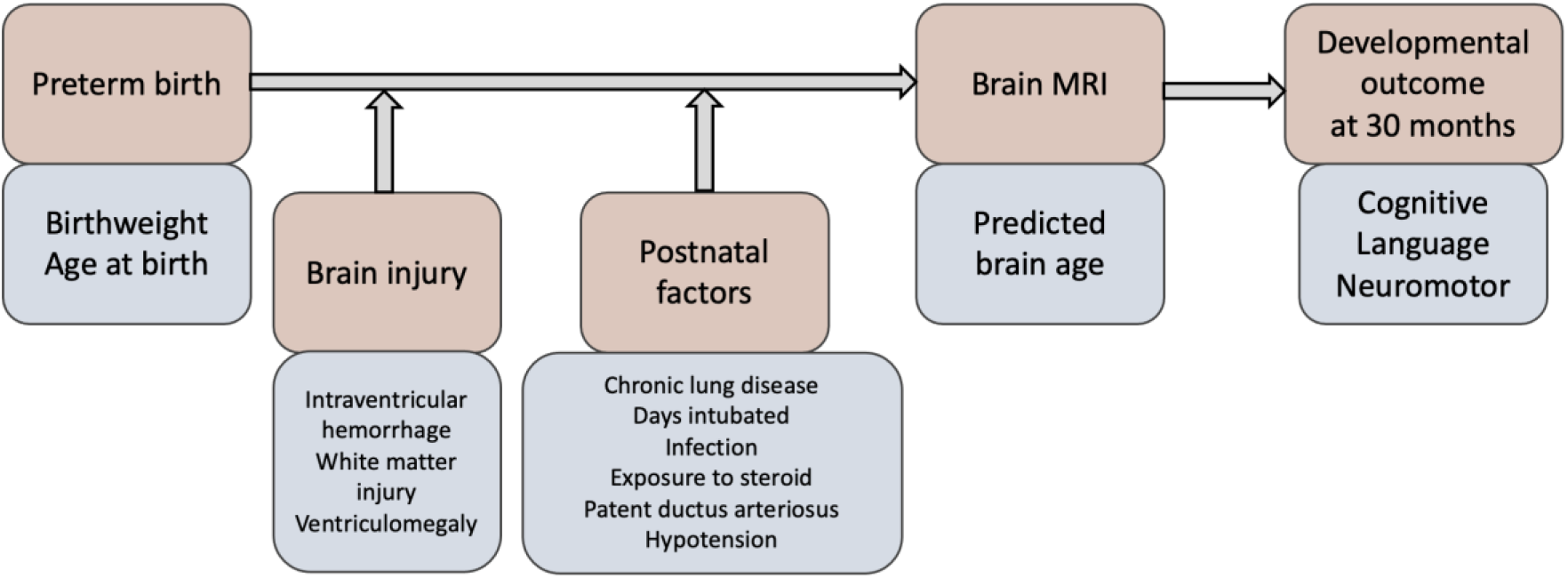
Hypothesized clinical risk factors and timeline after birth in relation to brain development in preterm infants. Orange: different stages when the given conditions, procedures, or measures (blue) occur.

By constructing the SEM, we tested our hypothesized model (Figure 2) using the aforementioned data and determined the strength and significance of each hypothesized path in our model. We estimated all parameters using weighted least squares with standard errors and mean- and variance-adjusted test statistics with a full weight matrix (WLSMV), which yields parameter estimates and standard errors that are robust to violations of multivariate normality (Power *et al.*, 2018). We used the χ^2^ statistic fit indices to evaluate whether the model fit to data well. We reported standardized parameter estimates computed using the WLSMV estimation to enable more direct comparisons of the effects for different pathways to neurodevelopmental outcomes.

To begin, we used the confirmatory factor analysis (CFA) to model latent variables summarizing a subset of pathologies. As a priori, we hypothesized a latent variable for 1) preterm birth measures, informed by birth age and birthweight; 2) perinatal injuries, informed by the presence of intraventricular hemorrhage (IVH), periventricular leukomalacia (PVL) or white matter injury (WMI), ventriculomegaly (VM); 3) postnatal conditions/treatments, informed by exposure to steroids (steroid), hypotension, infection, patent ductus arteriosus (PDA), days intubated (binarized), and chronic lung disease (CLD); 4) neurodevelopmental outcome scores in 30 months, informed by cognitive, language, and motor scores.

### Regional brain age prediction

We further applied the GCN for regional brain age prediction. To do this, the cortical surface was first parcellated into 76 cortical regions (ROIs) using the AAL atlas (Tzourio-Mazoyer *et al.*, 2002) that has been adapted and mapped onto the neonatal surface template. We used the original NOCIVET output surfaces that were extracted from each individual image with 81,924 vertices and partitioned these vertices and the related surface meshes into each ROI. Features mapped on the partitioned vertices together with the surface meshes as the topological graph within each cortical ROI (mean SD: 1,012 542 vertices per ROI) were fed to separate GCN models to predict the regional brain age (tested using 5-fold cross-validation). For each individual, we then computed an RBA per ROI.

To validate our hypothesis that RBA is associated with the brain development in future, we conducted linear regressions between each regional RBA and three neurodevelopmental scores at 30 months respectively. This analysis was performed on the 50 MR images of baseline and follow-up scans from 38 preterm neonates who revisited the center 30 months after their birth.

## Data availability

Anonymized data will be shared for reasonable requests from qualified investigators.

## Results

### Validation of GCN-based age prediction performance

We compared the prediction performance of each model on the entire dataset via 5-fold cross-validation. We found that the GCN fed with the true brain mesh achieved an accuracy of prediction with a mean absolute error (MAE) of 1.056 weeks (7.7% of the total age range) and absolute standard deviation (SDAE) of 0.943 weeks, which were smaller than the errors of the other two classic predictive models, i.e., random forest (RF) and general linear model (GLM) using the same number of vertices and features. Furthermore, to prove that the connections between vertices also contribute to the prediction of brain age, we conducted GCN-based prediction on randomized brain mesh structures (by disconnecting existing connections between random pairs of nodes and setting new connections between random pairs of nodes). Results indicated that the GCN model outperforms the other three models (Figure 3A, Table S2). Predicted brain age using GCN was also more accurate than the neonatal brain age predictions reported in the literature (MAE relative to total age range: 8.6-11.1%)(Brown *et al.*, 2017; Kawahara *et al.*, 2017). The brain age predicted using GCN strongly correlated with PMA (r = 0.87; Figure 3B).

**Figure 3.**
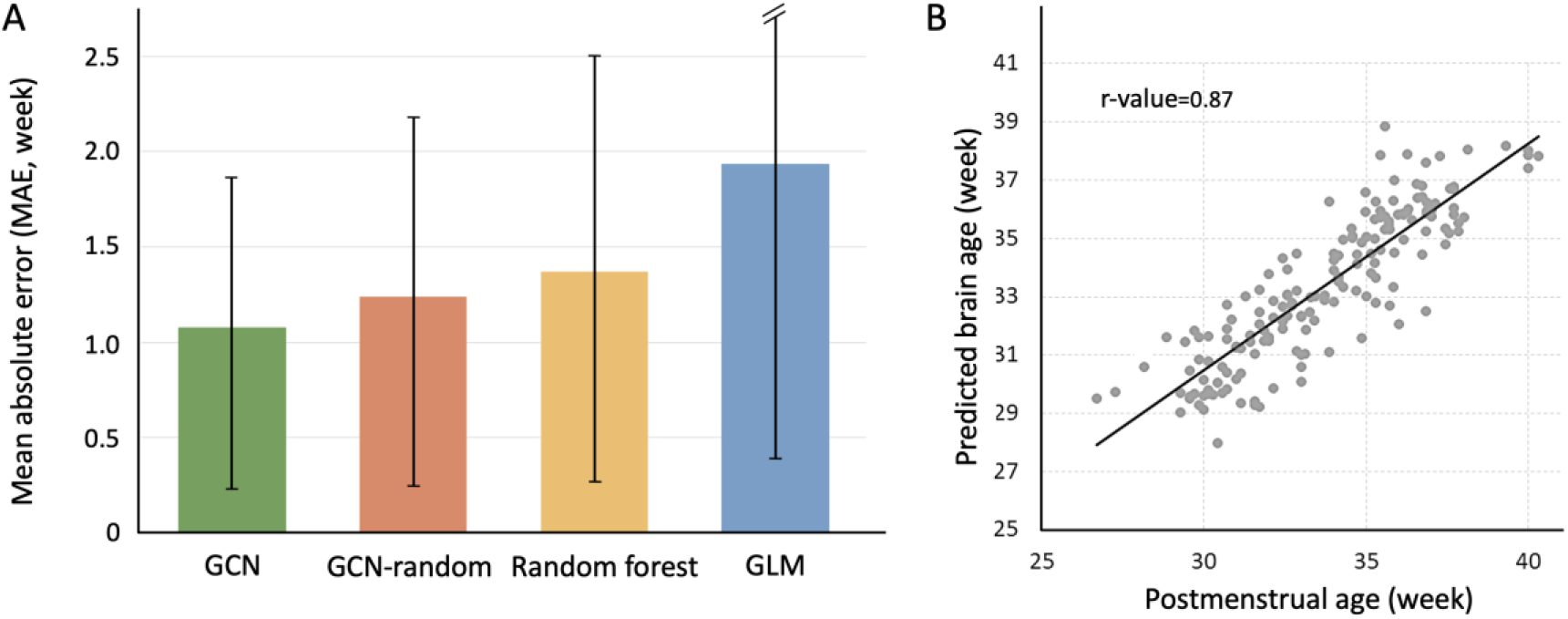
A) Brain age prediction error comparison among regression models. The height of each bar indicates mean MAE, and the black line indicates the standard deviation of MAE per model. GCN demonstrates the best prediction results. B) Scatter plot displaying brain age predicted using GCN model *vs.* chronological brain age.

To understand the relationship between cortical morphological features and the true age, we correlated the three features with PMA, and found that correlation coefficient r-values were 0.76 for cortical thickness, 0.80 for sulcal depth and −0.57 for GM/WM intensity ratio.

### RBA reflects the influence of perinatal clinical factors on brain growth in preterm neonates

We used relative brain age (RBA) measurements after bias correction described in the Methods. The univariate analysis showed that various clinical variables were associated with lower RBA (t>3.1; p<0.05, all p-values reported here and in the following sections were adjusted by the false discovery rate; Figure 4). Postnatal steroid exposure (p = 0.007) presented the strongest association with lower RBA. Chronic lung disease (CLD) (p = 0.031) and birthweight lower than 1000g (p = 0.021) were also significantly associated with lower RBA. These patterns were not found when using conventional RBA without bias correction (p’s > 0.05).

**Figure 4.**
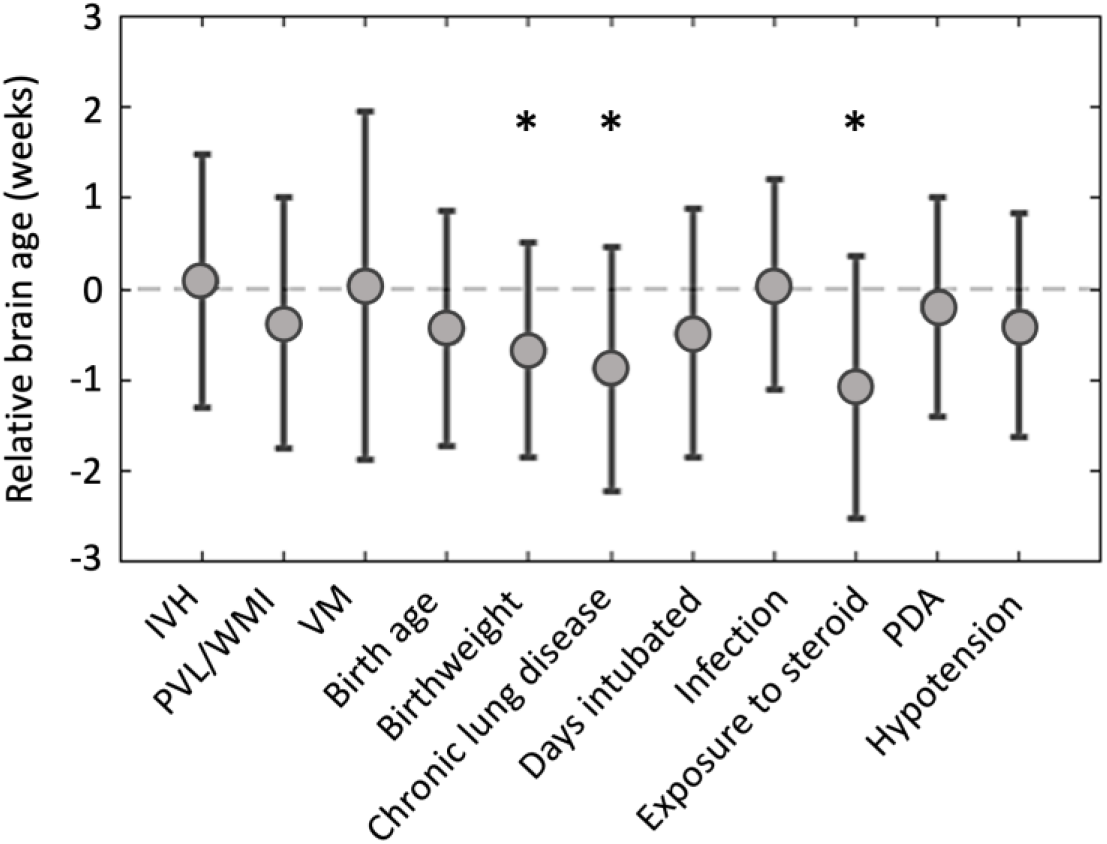
lower RBA representing impaired brain development was related to clinical factors.

### Relative brain age is associated with 30 months’ neurodevelopmental outcome in preterm neonates

Correlation of RBA with cognitive, language, or neuromotor scores in Bayley-III Scales of Toddler Development (Figure S3) showed that, after bias correction as in Figure 2, a lower RBA at neonatal scan was significantly associated with lower cognitive performance (r=0.41; p = 0.0025) and lower language performance (r=0.27; p = 0.0419). However, these patterns were not found when using conventional RBA without bias correction (p’s > 0.05).

### Association of regional brain age with clinical factors and neurodevelopmental outcome

We examined which brain regions predict the age at scan more accurately and whether specific regional RBAs are associated with either perinatal clinical factors or neurodevelopmental outcomes scored at 30 months of age.

Based on 5-fold cross-validation, age prediction for regional meshes resulted in a range of prediction accuracy with MAE of 1.19-2.42 weeks (Figure 5). We also found that the 9 smallest ROIs that contained less than 400 vertices (left, right olfactory cortices; left and right temporal poles of middle temporal gyrus; left and right Heschl gyri; left and right middle frontal gyri – orbital portion, right posterior cingulate cortex) displayed much larger MAEs (>1.75 weeks), possibly explaining the lack of fitting in the model due to a small number of features. Other 65 ROIs showed more reliable performance with MAEs less than 1.6 weeks. The pattern of prediction accuracy for cortical regions was hemispherically symmetric. The left middle cingulate cortex showed the most accurate prediction for NMI brains (MAE: 1.19 weeks), followed by left posterior cingulate cortex and right middle cingulate cortex (1.21 weeks), which were slightly larger than the prediction error computed using the whole brain surface data (1.06 weeks). These three regions also presented the highest slopes of brain age over the chronological age, which indicated their faster development.

**Figure 5.**
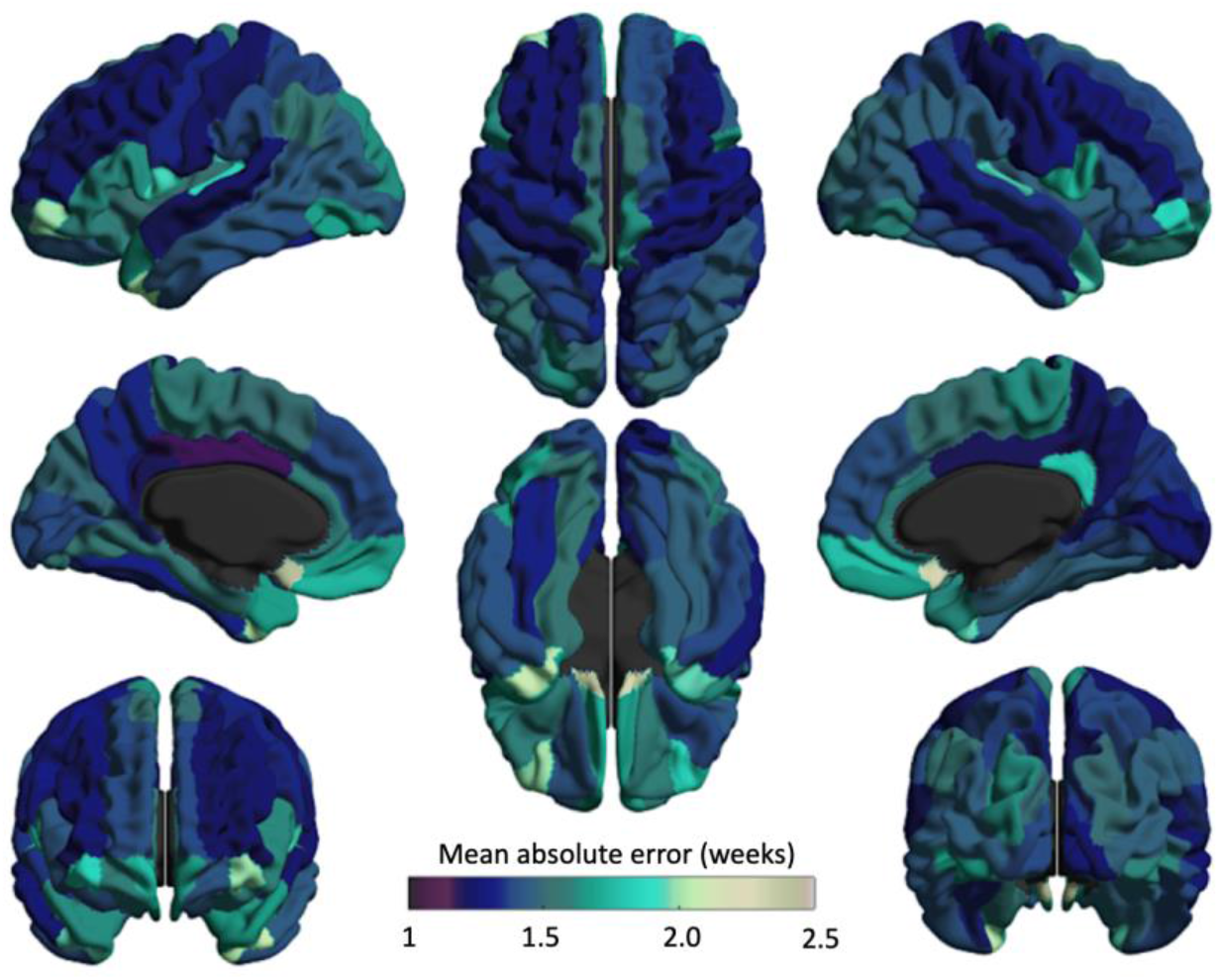
Mean absolute error map for regional RBAs predicted using GCN models.

We then performed the univariate analysis to assess the association between regional RBA and each of the clinical variables that showed significant correlations with the global RBA. Results after executing the Benjamini & Hochberg procedure for controlling the false discovery rate (FDR) showed that postnatal steroid treatment is associated with lower RBAs in five brain regions (t>3.1; p < 0.05), including left and right postcingulate cortices, right orbitofrontal cortex, left Rolandic operculum cortex, and left cuneus cortex (Figure 6A). In addition, lower birthweight (less than 1000g) was associated with lower RBAs (t>3.4; p < 0.05) in the left paracentral lobule and right postcingulate cortex.

**Figure 6.**
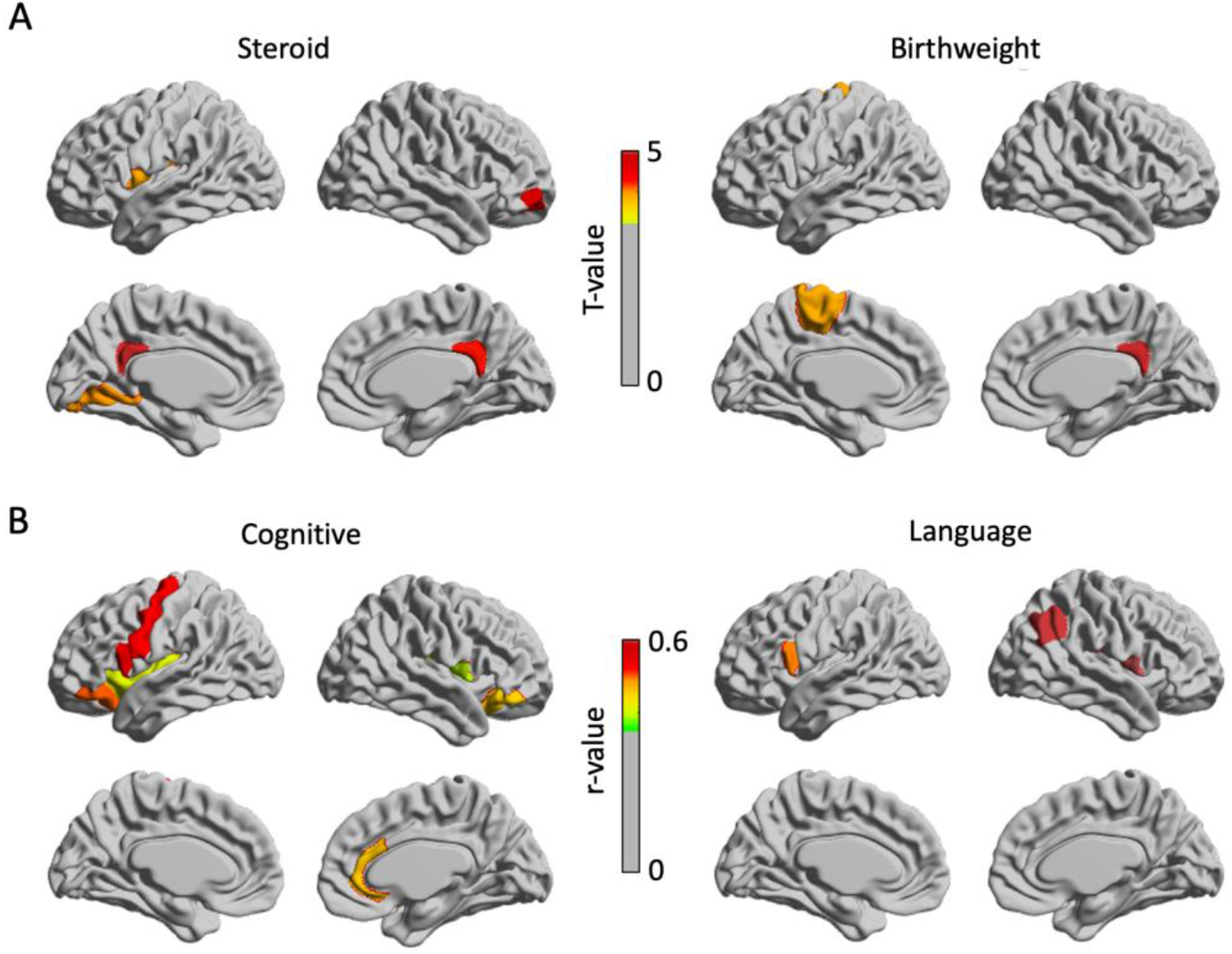
Association of Regional RBAs predicted using GCN models with perinatal clinical variables and neurodevelopmental outcome. **A.** Regional RBAs reflect the influence of perinatal clinical variables on brain growth. Left: brain regions displaying lower RBA associated with postnatal steroid treatment. Right: regions displaying lower RBA associated with extremely low birthweight (< 1000g). **B.** Regional RBAs are positively associated with brain cognitive (left), and language functional (right) scores at 30 months.

We then assessed correlations between regional RBAs and neurodevelopmental outcomes (Figure 6B). After FDR correction, six regions, including the left precentral, superior frontal, left inferior orbitofrontal, left insular, right Rolandic operculum right orbitofrontal, and right dorsal anterior cingulate cortices, displayed significant associations between their RBAs and cognitive scores at 30 months (r=0.41-0.53). Three regions, including left Broca’s area, right Rolandic operculum and right supramarginal cortex were significantly associated with language scores at 30 months (r= 0.46-0.57). No regional RBAs were associated with motor scores. No right hemispheric regional RBAs were associated with any domain of neurodevelopmental scales.

### Structural Equation Modeling

The outlined structural equation model is shown in Figure 7 (with standardized estimates), in which the rectangles represent observed variables, circles represent latent variables and error terms (e), single-headed arrows represent the impact of one variable on another, and double-headed arrows represent covariances between pairs of variables. The χ^2^ model-fit statistics indicated a significantly acceptable model fit (p < 0.001).

**Figure 7.**
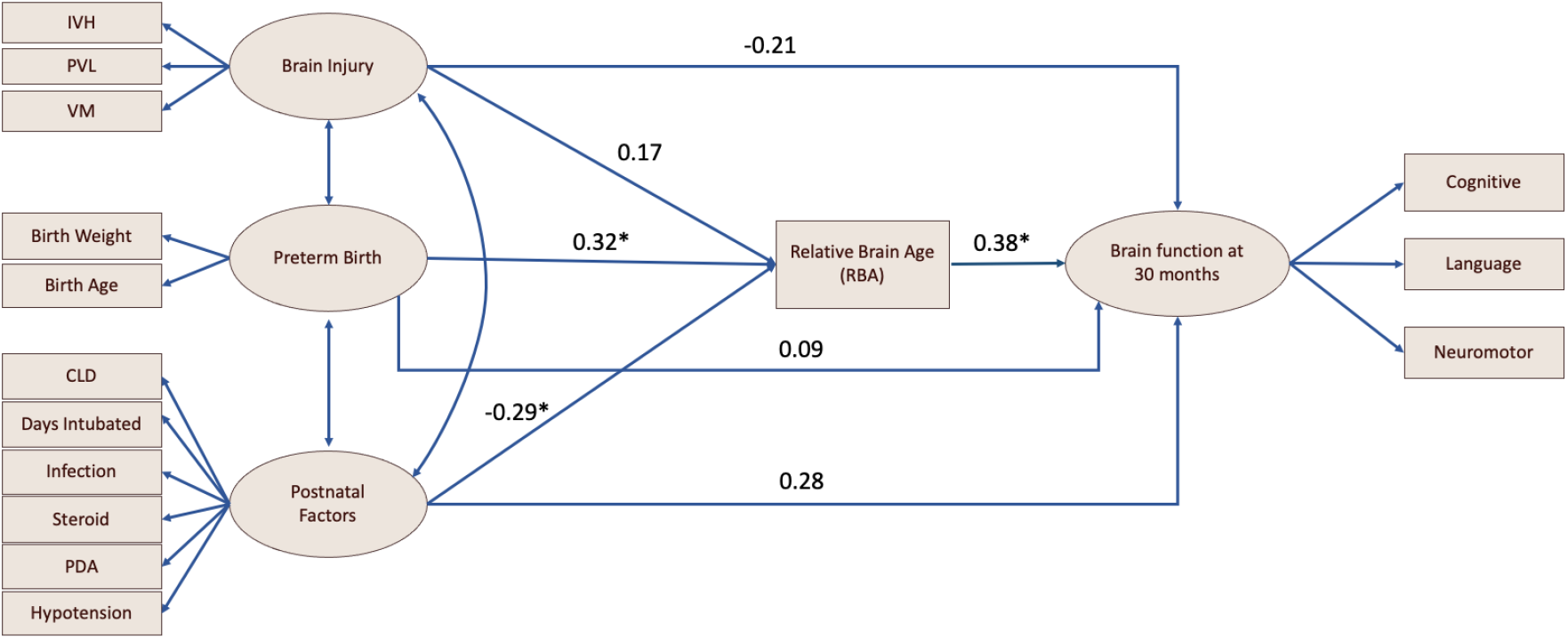
Results of path analysis. Rectangles represent manifest variables and ellipses represent latent variables. Each single-headed arrow denotes a hypothesized unidirectional effect of one variable on another. Numbers associated with effects are standardized regression coefficients. Asterisks refer to the paths that are statistically significant.

RBA was found to mediate the pathway from preterm birth to brain functional development at 30 months. RBA also mediates the pathway from postnatal factors to brain functional development at 30 months (p < 0.05). The relationship between brain injury and brain development at 30 months was not significant, for either the direct pathway or the pathway mediated by RBA (p < 0.05).

## Discussion

### Brain age prediction using GCN

With the capability of capturing the huge age-related morphological alterations during the third trimester, we proposed that GCN-based deep learning with surface morphological features can better predict individual brain age in this period with greater clinical utility in assessing neurodevelopmental status. To date, brain age prediction approaches have used machine learning methods fed with volume images directly (Cole *et al.*, 2017) or structural connectome metrics (Brown *et al.*, 2017). However, image-fed standard CNNs may not be capable of detecting the morphology that varies along the cortical manifold, which appears to be a sensitive gauge of brain growth in early development. Connectome-based deep learning approaches, such as the BrainNetCNN (Kawahara *et al.*, 2017), have also not performed better than conventional machine learning methods (He *et al.*, 2018).

To address these limitations, we incorporated morphological features extracted from the cortical surface and cortical surface topology into our GCN and found that its prediction accuracy is superior to state-of-the-art methods. We also demonstrated that learning of surface topological patterns is key to improving accurate prediction of the GCN fed with a surface of randomized connections, which have not been considered in other approaches. Using the GCN, we found that the morphological maturation of preterm neonatal brains can decelerate in the presence of various clinical conditions. The patterns found in regional age prediction indicate that the middle cingulate cortex is the best predictor for brain age in terms of the MAE. This finding is supported by previous evidence that confirms the cingulate area as one of the earliest developing regions and the thickest part of the cortex in newborns (Li *et al.*, 2015b).

### Clinical Implications and Applications

Our findings suggest that RBA has potential clinical utility in assessing neurodevelopment of preterm infants. Specifically, it is the first time using structural equation modelling (SEM) analysis to provide clear evidence of the temporal relationship among three stages during the developmental trajectory for neonates: the context the preterm neonates were placed in, the brain development and the behavioral patterns years later. That is, RBA at scan mediated the pathway from preterm birth and postnatal factors, such as exposure to postnatal steroids, CLD, hypotension, PDA, infection, days for intubation, to brain functional development at 30 months (Fig. 7). In addition, the significant pathway from preterm birth and postnatal factors to brain functional development at 30 months were not found. This suggests that brain morphological growth affected by preterm birth and postnatal factors is key to understanding neurodevelopmental impairment presented in preterm children. SEM did not provide sufficient evidence of a significant relationship between perinatal brain injuries and brain development at 30 months. Despite a limited sample size used for the analysis, these findings suggest that the effects of preterm birth and other postnatal factors on brain functional development may be larger than the effects of preterm birth-related brain injuries.

Given a close look at the association between single clinical factor and RBA, our GLM analysis showed that a lower RBA value, indicating delayed neurodevelopment, is significantly associated with several clinical factors, such as birthweight, CLD, and use of postnatal steroids (Fig. 4 and 6). The underlying pathophysiology of how these clinical factors impair neurodevelopment may be attributed to both exogenous and endogenous factors that lead to brain insult from respiratory or circulatory insufficiency, hemorrhagic events, hypoxic-ischemic events, and inflammation (Dempsey and Barrington, 2007; van Vliet *et al.*, 2013; Lemmers *et al.*, 2016; Galinsky *et al.*, 2018; Zonnenberg *et al.*, 2019).

Among the clinical variables analyzed, birthweight, CLD, and steroids demonstrated the most significant relationships with RBA impairment. Specifically, lower birthweight was associated with impaired RBAs of two specific brain regions, including the left paracentral lobe and right post-cingulate cortex (Fig. 6). Reduced RBAs in these regions implicate lower birthweight’s adverse impact on related functions of the paracentral lobule (i.e., motor and sensory functions of the lower limb as well as autonomic functions) and diverse functions of the right post-cingulate cortex (i.e., communication with brain networks, working memory, and the default mode network). Impairment of these regions are consistent with prior studies establishing associations between low birthweight infants and poor neurodevelopmental outcomes, such as developmental delay, movement disorders, visual problems, hearing impairment, and intellectual disabilities (de Kieviet *et al.*, 2009).

Our results also demonstrated that CLD has a strong association with impaired RBA. CLD primarily affects preterm infants who are exposed to prolonged mechanical ventilation and oxygen therapy for pulmonary complications (Kinsella *et al.*, 2006; Gallini *et al.*, 2020). An immature lung with chronic exposure to ventilation can result in oxygen toxicity and pulmonary inflammation, compromising proper gas exchange and thus perfusion and oxygenation to the brain. Consequently, the immature brain, which is vulnerable to these fluctuations in perfusion and oxygenation, becomes susceptible to a cascade of complications, including hypoxia-ischemia, inflammation, germinal matrix injury, diffuse white matter injury, and diffuse gray matter injury (Albertine, 2012; Malavolti *et al.*, 2018). These resulting complications, particularly diffuse white and gray matter lesions, contribute to neurodevelopmental delay in motor skills, learning, attention, and behavior (Perlman, 2001), consistent with the poor neurodevelopmental outcomes of CLD infants, including motor (Van Marter *et al.*, 2011), cognitive (Singer *et al.*, 2001), and language deficits (Singer *et al.*, 1997; Short *et al.*, 2003; Natarajan *et al.*, 2012). Thus, our findings further support that systemic complications of CLD contribute to neurodevelopmental delay as indicated by its strong association with reduced RBA.

Furthermore, our results revealed that postnatal exposure to steroids have an association with RBA impairment. Specifically, use of postnatal steroids impaired RBAs of five brain regions, including the left and right post cingulate cortex, right orbito-frontal cortex, left Rolandic operculum cortex, and left cuneus cortex (Fig. 6). Postnatal steroid therapy is traditionally provided to preterm neonates with BPD to reduce lung inflammation (Halliday *et al.*, 2003). In our previous study, postnatal exposure to clinically routine doses of hydrocortisone or dexamethasone were associated with impaired cerebellar but not cerebral growth when total volumes were analyzed (Tam *et al.*, 2011). Using RBA based on cortical morphometrics and deep learning algorithms, however, we revealed adverse effects of postnatal glucocorticoids on cerebral growth. These results are consistent with previous findings that poor neurofunctional development following postnatal steroid usage in other studies (Watterberg, 2010; Cheong and Doyle, 2019; Van Meurs and Hintz, 2020).

Finally, the potential clinical utility of RBAs is further supported by their strong relationships with cognitive and language scores at 30 months (Fig. 6B). Specifically, cognitive scores were significantly associated with RBAs of six brain regions, including the left precentral, superior frontal, left inferior orbitofrontal, left insular, right Rolandic operculum, right orbitofrontal, and right dorsal anterior cingulate cortices. Furthermore, language scores were significantly associated with RBAs of three brain regions, including left Broca’s area (important for language production and comprehension), right Rolandic operculum, and right supramarginal cortex. These brain regions are involved in specific functions of cognition and language process: (1) the post cingulate cortex important for memory, emotional regulation, the default mode network, and communication with other networks, (2) the orbitofrontal cortex important for cognitive processing and decision-making, (3) the Rolandic operculum cortex important for somatosensory and motor function, and (4) the cuneus important for visual processing. These findings show a broad overlap between affected brain regions and their respective functional roles in cognition and language, further supporting the potential of RBAs in accurately predicting neurofunctional development. Altogether, our findings are largely consistent with existing evidence revealing adverse effects of prematurity-related clinical variables on the developing brain’s structure and function (Shah *et al.*, 2008; Brew *et al.*, 2014; Kidokoro *et al.*, 2014; Lemmers *et al.*, 2016; Van Meurs and Hintz, 2020).

We found that our RBA metric was not associated with neuromotor scores at 30 months. A possible explanation is that the morphology in the motor cortex, particularly cortical folding, was already relatively mature in our cohort, given folding forms earlier than the late 3^rd^ trimester of gestation. Therefore, predicted brain age based on cortical morphology extracted from these postnatal scans may not be sensitive to neuromotor impairment, while previous studies have shown that neuromotor impairment of preterm survivors is rather significantly associated with perinatal white matter brain injury (Guo *et al.*, 2017; Saha *et al.*, 2020) and postnatal cerebellar growth (Messerschmidt *et al.*, 2008).

### Limitations, future directions and conclusions

The acceleration/deceleration of predicted brain age could be a surrogate for brain developmental status, which may reflect a combination of perinatal clinical factors exert on preterm neonates before the scan and be associated with brain functional abilities in the future. However, the *conventional RBA metric is not suitable for gauging the degree of brain health* due to its bias. We further showed that this metric fails to show significant difference for any of the clinical factors and predict preterm survivors’ functional outcomes. By correcting this bias, we demonstrated that the adjusted RBA measurement showed significant difference for some of the clinical conditions, also it significantly correlates with cognitive and language scores evaluated after 30 months. It is worth noting that the relative brain age is frequently influenced by various sources that we could not consider fully, which may give rise to significant false positives and false negatives when looking for associations between RBA and other measures (Le *et al.*, 2018). In this study, we only corrected the linear bias from the relative brain age. Suggested by Smith et al. (Smith *et al.*, 2019), it is necessary to not only remove the linear dependency of relative brain age on age but also the nonlinear dependence, especially the quadratic dependency of brain aging (as a function of age).

There are several other limitations of the current study. Due to the modest sample size and high-dimensional feature space, it is possible that some of the reported effect sizes are over-fitted. Future studies should focus on replicating the current findings using a larger sample size. To better investigate the relative brain age in regard to injuries and clinical factors, it is best to have brain age prediction models trained through healthy control subjects. However, our preterm neonate dataset includes only a few completely “healthy” subjects displaying no clinical conditions: i.e., 7 out of 170 neonates. Hence, our brain age prediction model in this current study was not trained solely based on “healthy” preterm neonates who presented no clinical conditions, which contributes to a degree of bias from existing clinical variables.

We found that very small cortical regions displayed much larger error in prediction of brain age, possibly explained by the lack of morphological features used for the fitting process. Yet, it is noteworthy to mention that there were significant associations between the RBA of two of those small regions (right posterior cingulate, right orbital portions of middle frontal gyrus) and perinatal risk factors, suggesting the importance of these small regions in neonatal development under specific conditions. To fully clarify, these cortical regions remain to be analyzed more carefully with a larger sample dataset.

Despite these limitations, however, our study proposes a novel GCN that uniquely uses the dramatically altered morphological features and topological patterns to predict brain age, which in turn explain the developmental trajectory in preterm neonates by linking to pre-scan clinical factors and post-scan behavioral developments. Our study contributes to the growing body of literature on early postnatal neurodevelopment. Altogether, these findings provide the basis for future investigations aiming to extend the PBA measurement to practical clinical application, such as the individualized prediction of neurodevelopmental outcome alongside other potential biomarkers of brain development.

