## supplementary for "Brain age predicted using graph convolutional neural network explains developmental trajectory in preterm neonates"

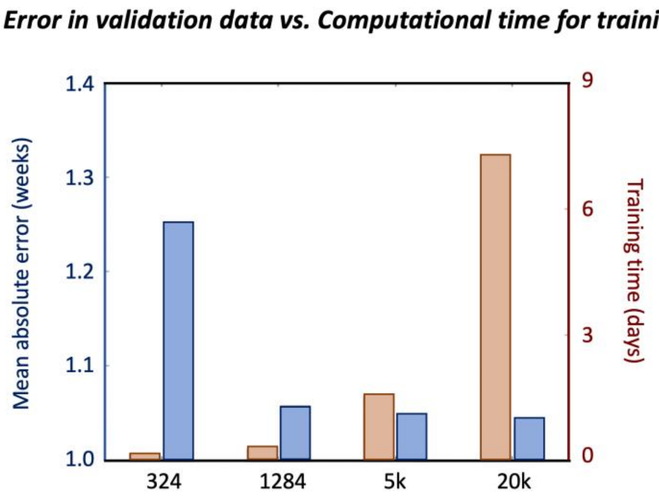

8

Figure S1. The number of 1,284 was chosen to use in the following analysis by a compromise of the computational accuracy and computational time

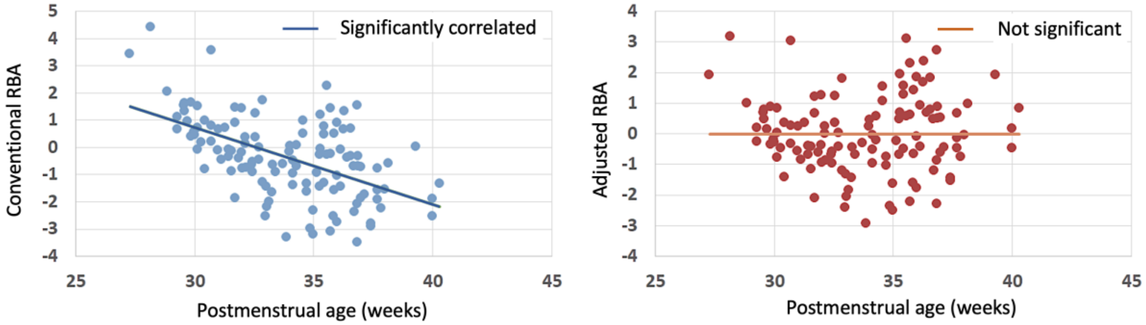

Figure S2. The conventional RBA is negatively associated with true age (left), while this effect is corrected in new RBA (right).

Table S1. Definition and categorization of clinical variables for statistical analysis.

| Factors | Definition | Grouping | |
|---|---|---|---|
| IVH | Papilie et al. 1978 | grade 0-2 (143) | 3-4 (25) |
| VM | Miller et al. 2003 | grade 0-1 (114) | 2-3 (18) |
| PVL (WMI) | Miller et al. 2003 | grade 0-1 (152) | 2-3 (56) |
| Birth_Weight | gram | >=1000 (72) | 1-999 (93) |
| Birth_Age | weeks | >=28 (83) | <28 (87) |
| Days_Intubated | Mechanical ventilation and intubation | 0-5 (87) | >=5 (57) |
| PDA | Patent ductus asteriosus | No (69) | Yes (87) |
| CLD | | No (107) | Yes (74) |
| Neonatal_Infection | Culture + or – sepsis, and/or meningtis | No (81) | Yes (89) |
| Postnatal_Steroid | Exposure to postnatal hydrocortisol | No (134) | Yes (21) |
| Hypotension | Requiring treatment with volume | No (50) | Yes (103) |

| | resuscitation and/ or vesopressors required |
|---|---|

Table S2. The performance of the age prediction (errors in weeks for MSE, MAE and SDAE).

| | MSE | MAE | SDAE | r-value |
|---|---|---|---|---|
| Cortical thickness | 2.501 | 1.991 | 1.682 | 0.60 |
| Sulcal depth | 2.477 | 1.982 | 1.675 | 0.62 |
| GM/WM intensity ratio | 2.889 | 2.372 | 1.775 | 0.39 |
| General linear model | 2.432 | 1.938 | 1.655 | 0.70 |
| Random forest regression | 1.796 | 1.373 | 1.165 | 0.81 |
| GCN with random mesh | 1.593 | 1.239 | 1.027 | 0.85 |
| GCN with true mesh | **1.499** | **1.056** | **0.943** | **0.87** |

Abbreviation: MSE = mean squared error; MAE = mean absolute error; SDAE= absolute standard deviation

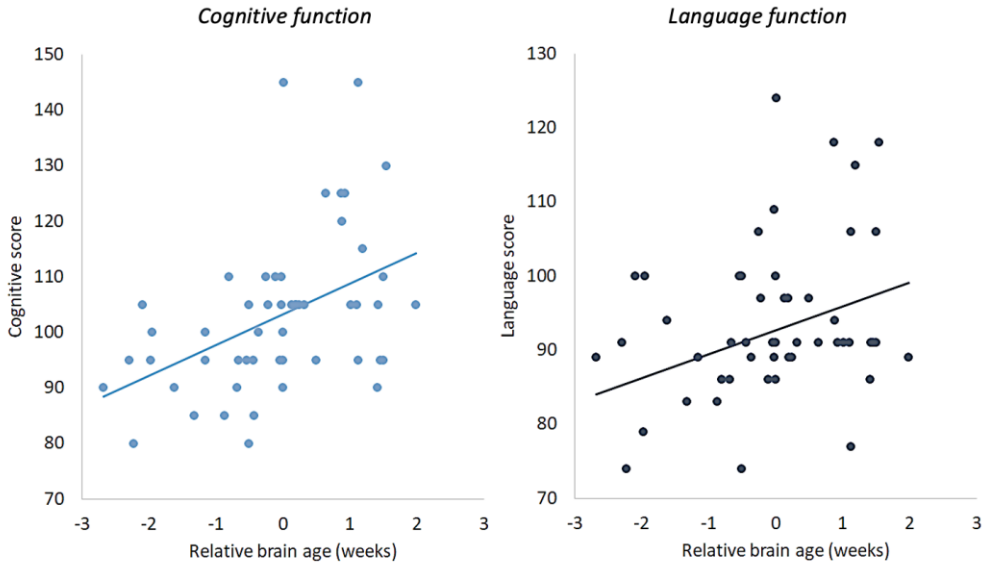

Figure S3. Correlation between RBA and neurodevelopmental outcomes at 30 months (left: cognitive score; right: language score).
